# NMR studies reveal that protein dynamics critically mediate aggregation of the well-folded and very soluble *E. coli* S1 ribosomal protein

**DOI:** 10.1101/178459

**Authors:** Yimei Lu, Liangzhong Lim, Jianxing Song

## Abstract

Unlike mammalian aging associated with many hallmarks, *E. coli* aging is only significantly characterized by protein aggregation, thus offering an excellent model for addressing the relationship between protein aggregation and aging. Here we characterized conformations, unfolding and dynamics of ribosomal protein S1 and its D3/D5 domains using NMR, CD and fluorescence spectroscopy. S1 is a 557-residue modular protein containing six S1 motifs. Paradoxically, while S1 is well-folded and very soluble *in vitro*, it was found in various lists of aggregated *E. coli* proteins. Our results decipher: 1) S1 has dynamic inter-domain interactions. Strikingly, S1 and its D3/D5 domains have significantly exposed hydrophobic patches characterized by irreversible unfolding. 2) Although D5 has significantly restricted backbone motion on ps-ns time scale, it has global μs-ms conformational dynamics and particularly high “global breathing” motions. 3) D5 assumes the conserved β-barrel fold but contains large hydrophobic patches at least dynamically accessible. Taken together, our study reveals that S1 could be prone to aggregation due to significant dynamics at two levels: inter-domain interactions and individual domains, which may even render buried hydrophobic patches/cores accessible for driving aggregation. This mechanism is most likely to operate in many proteins of *E. coli* and other organisms including human.

---

Protein aggregation was not only characterized to be problematic for *in vitro* protein research and industry applications, but has been also identified to commonly lead to a large spectrum of human diseases^1–12^, which include all human neurodegenerative diseases^1–6^, such as Parkinson’s disease (PD), Alzheimer’s disease (AD), Huntington’s disease (HD), spinocerebellar ataxias (SCA), amyotrophic lateral sclerosis (ALS); as well as diabetes^7^ and cardiac dysfunction^9^. Furthermore, aggregation of a large number of non-specific proteins particularly overrepresented by the β-architecture proteins has been shown to be associated with aging of all organisms^12–18^. Most intriguingly, asymmetric segregation of protein aggregates has been recently revealed to be characteristic of the cellular aging and rejuvenation of *E. coli* cells^16,17^.

Aggregation of specific proteins has been demonstrated to lead to human diseases by “loss of functions” or/and “gain of toxic functions”. However, as even for the human neurodegenerative diseases, the involved proteins are so functionally and structurally diverse and their knockout (such as SOD1) does not always result in the corresponding diseases. Consequently it is most likely there is no common mechanism underlying “loss of functions”. On the other hand, proteins involved in all these diseases are universally prone to aggregation and therefore there should be a common mechanism for “gain of toxic functions”. As such, to decipher the molecular mechanisms underlying aggregation of associated proteins represents a central focus as well as a key step in further delineating the mechanisms for “gain of toxic functions” to cause diseases.

By contrast, the mechanisms for aggregation of a large amount of non-specific proteins during aging have been less investigated and whether the aggregated proteins commonly gain toxic functions responsible for aging remains largely unknown. In the present study, we aimed to explore the mechanisms of aggregation of *E. coli* proteins because asymmetric segregation of protein aggregates appears to be the only significant biomarker characteristic of its cellular aging and rejuvenation^16,17^. Here we decided to select a cytosolic protein which satisfies two criteria: 1) the protein should be extensively found in various lists of *E. coli* aggregated proteins. 2) It is composed of well-structured folds rich in β-sheets because aggregated proteins associated with aging have been previously shown to be overrepresented by the β-dominant proteins^12–15^. Consequently, *E. coli* S1 ribosomal protein was selected as it was identified to aggregate severely in an *E. coli* strain carrying the mutant of YajL protein^19^, a prokaryotic homolog of parkinsonism-associated protein DJ-1 which functions to repair proteins in response to the global stress, particularly oxidative stress associated with aging^19–22^. Furthermore, the aggregate of S1 ribosomal protein was even identified in healthy *E. coli* cells^23^. Also paradoxically, the S1 ribosomal protein has been previously characterized to be well-folded and very soluble *in vitro* with the concentration even reaching up to 20-30 mg/ml^24,25^.

*E. coli* ribosomal protein S1 is the largest protein of eubacterial ribosomes^24–31^, which is a 557-residue modular protein composed of six independently-folded repeats of the S1 motif (Fig. 1A), a structural motif commonly found in proteins involved in RNA metabolism. Each repeat consists of about 70 amino acids separated by spacers of ∼10 residues, which is arranged into a five-stranded antiparallel β-barrel resembling the fold of the bacterial cold shock protein (Fig. 1B and S1)^32^. The first two repeats, D1 and D2, constitute the N-terminal 30S ribosome binding domains, while the rest of four repeats D3-D6 are involved in mRNA recognition. Due to its functional importance, previously ribosomal protein S1 has been extensively investigated by various biophysical methods^24–31^. However, the full-length S1 protein was not amenable to structure determination by X-ray crystallography in both free state and in complex with ribosome. Nevertheless, all six S1 modular repeats have been previously investigated by NMR spectroscopy and the atomic resolution structures have been determined in solution for the domains 1, 2, 4 and 6 (Fig. S1)^28–30^. Intriguingly, although all four structures have the conserved β-barrel, the lengths and orientations of the β-strands are very diverse. Furthermore, four structures have very variable helical segments (Fig. S1), and consequently it is almost impossible to structurally overlay four structures together.

**Figure 1.**
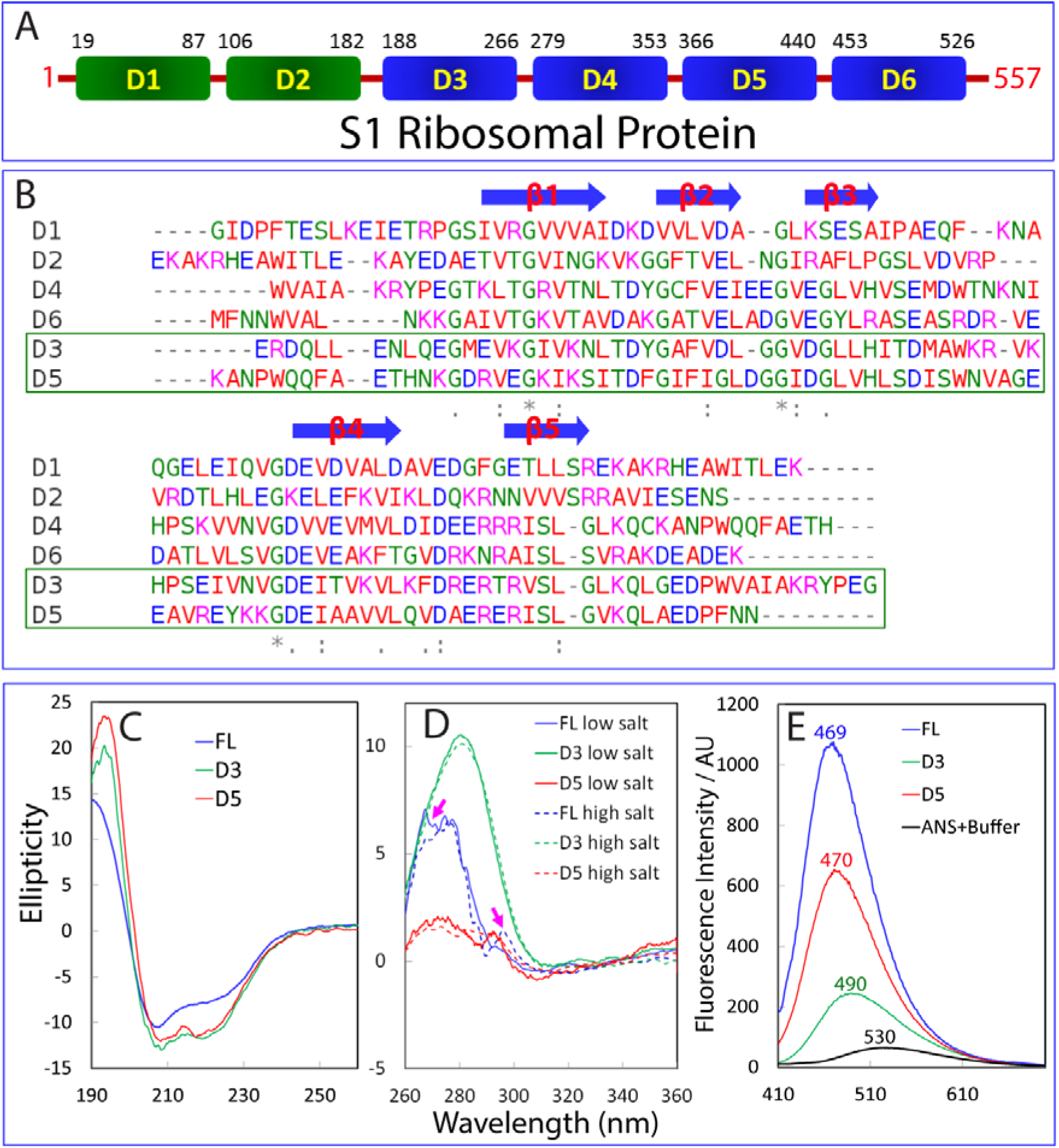
Organization and conformations of *E. coli.* ribosomal protein S1 and its D3/D5 domains. (A) *E. coli.* ribosomal protein S1 which is a 557-residue modular protein composed of six repeats of the S1 motif. The first two domains, D1 and D2 (green), constitute the N-terminal 30S ribosome binding domains, while the rest of repeats D3-D6 (blue) are responsible for mRNA recognition. (B) Alignment of six S1 repeats with the sequences of the D3 and D5 domains we studied here boxed. Five conserved β-strands are indicated by blue arrows. (C) Far-UV CD spectra of the full-length S1 protein and its D3 and D5 domains collected in 1-mm curvet at 25 °C at a protein concentration of 15 μM in the low salt buffer. (D) Near-UV CD spectra of the full-length S1 protein and its D3 and D5 domains collected in 10-mm curvet at 25 °C at a protein concentration of 50 μM in the low salt buffer (solid lines); and in the high salt buffer (dashed lines). The purple arrows are used to indicate the differences in the near-UV signals of the full-length S1 protein in low and high salt buffers. (E) ANS fluorescence spectra of the full-length S1 protein and its D3 and D5 domains at a protein concentration of 20 μM in the presence of 80 μM ANS in the high salt buffer with 2 mM DTT. The fluorescence intensity was reported in arbitrary unit.

While extensive studies have been previously carried out to delineate the structure-function relationship of the protein S1 and its isolated domains, the mechanisms underlying the aggregation remain largely unexplored by high-resolution biophysical methods. The full-length protein S1 has been previously found to be highly soluble *in vitro* in relatively low-salt buffers^24–27^, and its isolated D1, D2, D4 and D6 domains should be also highly soluble in order to allow previous NMR structure determinations, which requested stable protein samples at high protein concentrations^28–30^. Therefore, here we focused on characterizing solution conformations, thermal unfolding and dynamics of the full-length S1 proteins and its D3 and D5 domains by NMR, CD and fluorescence spectroscopy. The results reveal that consistent with the recent results^27,28^, the six well-folded S1 repeats are not independently behaved in the full-length S1 protein, but instead have dynamic inter-domain interactions. Furthermore, the S1 protein appears to have significant exposure of hydrophobic patches and consequently is prone to aggregation at high salt concentrations, or/and at high temperatures. Moreover, the isolated D3 and D5 domains also have significantly exposed hydrophobic patches and are characterized by the irreversible unfolding despite containing no Cys residue. Residue-specific NMR dynamic studies reveal that while the D5 domain has significantly restricted backbone motions on ps-ns time scale, it has global μs-ms conformational dynamics. In particular, both D3 and D5 domains have extremely high conformational dynamics on min-hr time scale. Taken together, the results reveal the critical roles of protein dynamics in aggregation of *E. coli* ribosomal protein S1, thus rationalizing the paradoxical observation that while the S1 protein is both well-folded and highly soluble, it could be prone to aggregation in cells with high salt concentrations.

## Results

### Solution conformations of ribosomal protein S1 and its D3 and D5 domains

Previously the overall shape of the ribosomal S1 protein has been extensively characterized by various biophysical methods^24–27^. Interestingly, while early studies indicated that it is a strongly asymmetric 10:1 prolate ellipsoid with a major axis 200-250 Å in length^24–26^, recent results suggested a compact globular conformation with dynamic inter-domain interactions^27^. Moreover, the solution structures of individual D1, D2, D4 and D6 domains have been determined to adopt the conserved S1 motif by NMR spectroscopy^28–31^. On the other hand, although the D3 and D5 domains have been previously characterized by NMR in the isolated domains as well as in the contexts of linking with other domains^28^, their NMR structures remain unsolved so far. Interestingly, previous results revealed that the isolated D3 domain formed a non-native dimer and the residues 260-277 to link D3 and D4 appeared to have extensive interactions with the D3 domain^28^.

Here, here we focused on characterizing the full-length S1 protein and its D3 and D5 domains. As compared with the D3 (181-266) and D5 (350-440) domains previously investigated^28^, our D3 construct consists of residues 181-277, thus covering the whole linker between D3 and D4 domain. Our D5 construct has the residues 350-443, with three extra residues at the C-terminus. While the full-length S1 and D3 domain were highly soluble in supernatant as previously reported^28^, our D5 construct was found to be partly soluble, with a portion of the recombinant D5 protein in supernatant if induced by a low IPTG concentration (0.3 mM) and at low temperature (20 °C), thus overcoming the previous problem that the D5 domain was completely found in inclusion body and thus needed to be purified under denaturing conditions. As such, here all three proteins have been successfully purified by Ni^2+^-affinity column under native conditions followed by further FPLC purification.

As shown in Fig. 1C, the full-length S1 protein has a far-UV CD spectrum very similar to that of the S1 protein directly purified from *E. coli* cells without over-expression^24^. The D3 and D5 domains have very similar CD spectra typical of a folded protein, consistent with the previous studies^28^. Furthermore, they also have tertiary packing as reflected by the presence of signals in the near-UV spectra (Fig. 1D)^33,34^. Interestingly, while the isolated D3 and D5 domains have very similar near-UV CD spectra in both low salt buffer (1 mM phosphate at pH 6.8), and high salt buffer (50 mM phosphate at pH 6.8 with 200 mM NaCl) which was previously used to mimic cellular conditions for NMR characterizing D3 and D5 domains^28^, the full-length S1 protein has slightly different near-UV CD spectra in two buffers, implying that salt ions may mediate its inter-domain interactions as previously revealed by other biophysical methods^27^.

To assess their surface properties, we measured their binding to a fluorescent hydrophobic probe 1-anilinonaphthalene-8-sulfonate (ANS) which has been widely used to measure exposed hydrophobic patches on proteins^35,36^. As shown in Fig. 1E, upon adding proteins, the emission maximum of ANS shifted from 530 nm to 469 nm for the full-length S1, to 490 nm for D3; and to 470 nm for D5, and the fluorescence intensity also significantly increased. The results indicate that the full-length S1 protein and its dissected D3/D5 domains all have significantly exposed hydrophobic patches^35,36^. Noticeably, out of three proteins, the full-length protein appears to have the highest amount of exposed hydrophobic patches.

We subsequently collected ^1^H one-dimensional (1D) and ^1^H-^15^N two-dimensional (2D) HSQC spectra for the full-length S1 protein (Fig. 2A). Although several very up-field 1D peaks characteristic of a folded protein could be observed, they are very broad as previously reported^25^. Consistent with 1D NMR spectrum, only a small number of HSQC peaks could be detected, which are all narrowly-dispersed and thus likely from the flexible and disordered regions. Furthermore, we conducted dynamic light scattering (DLS) measurements which indicate that if based on the assumption that the S1 protein is a tightly packed globular protein, its apparent molecular weight would be ∼270 kDa at protein concentrations of 100 and 300 μM, which is much larger than its theoretic molecular weight of ∼62 kDa. Taken together, the results implies that the six well-folded S1 repeats are not independently behaved with the completely-flexible linkers but instead these repeats have dynamic inter-domain interactions, completely consistent with recent studies by various biophysical methods^27,28^. As a result, most HSQC peaks from well-folded S1 repeats are undetectable due to a slow global tumbling resulting from the large size, or additional μs-ms dynamics. HSQC peaks could be detected only for the very flexible regions due to the “motional narrowing” effect^37,38^.

**Figure 2.**
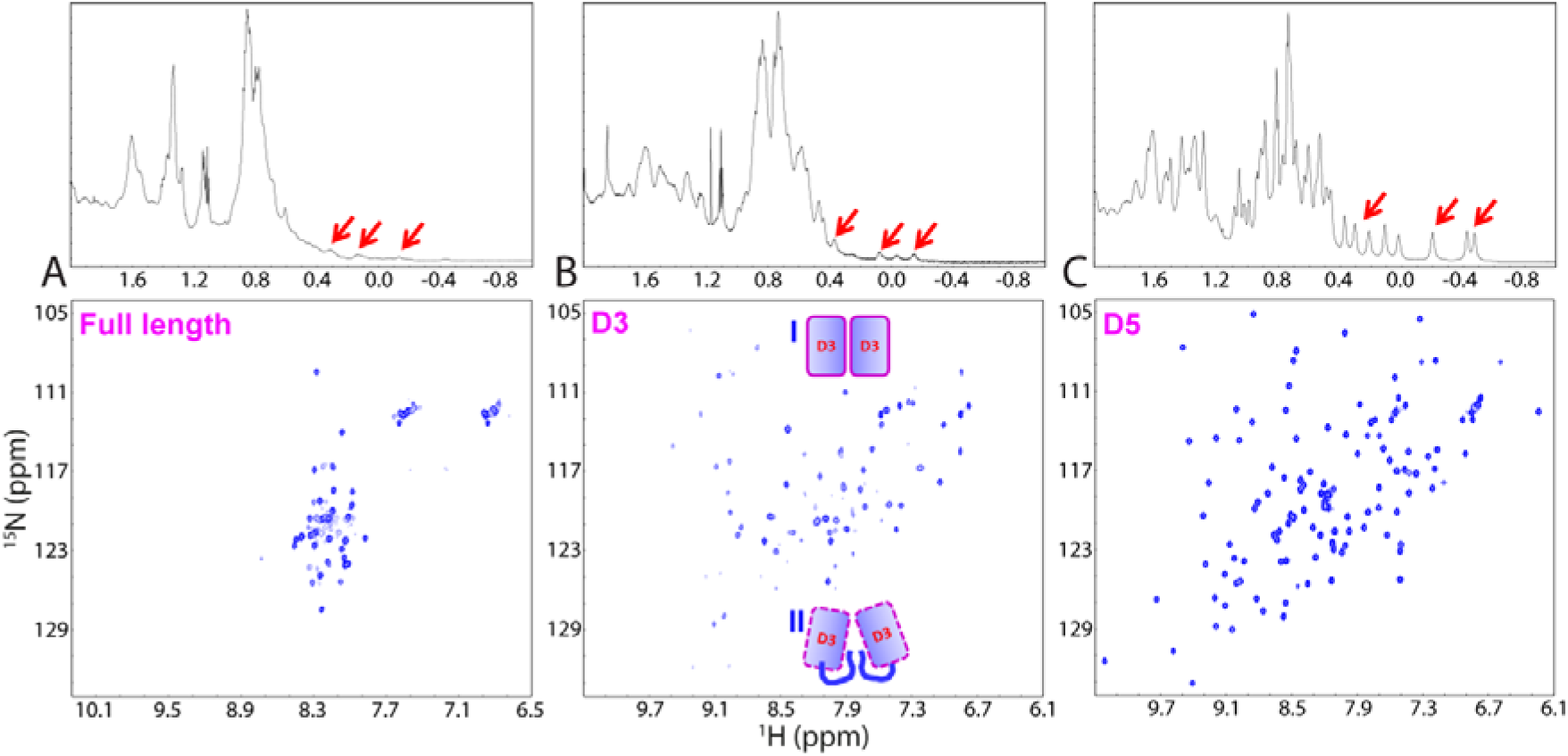
NMR characterization of S1 and its D3/D5 domains. One-dimensional (1D) ^1^H and two-dimensional (2D) ^1^H-^15^N HSQC spectra of the full-length S1 protein (A); D3 (B) and D5 (C) domains collected at 25 °C. Inlets of (B): model of the dimeric D3 (181-266) (I) previously investigated^28^, and D3 (181-277) (II) studied here. Red arrows are used to indicate very up-field 1D NMR peaks typical of a well-folded protein.

Interestingly, although our D3 (181-277) construct does have very up-field 1D NMR peaks typical of a folded protein, they are much more broad than those expected for a protein of ∼12 kDa (Fig. 2B). Consistent with significant line broadening of 1D peaks, the number of detectable HSQC peaks of the D3 domain (181-277) is only ∼80% of what was predicted from its sequence. This is completely different from what previously reported for D3 (181-266) that despite forming a non-native dimer with an apparent molecular weight of 23 kDa, it had a well-dispersed HSQC spectrum with almost all HSQC peaks for non-Pro residues detectable and assigned^28^. DLS measurements indicate that our D3 domain has an apparent molecular weight of ∼31.4 kDa at protein concentrations of 100 and 500 μM, even larger than the dimeric form of the previous D3 construct^28^. As such, it appears that as compared to the short D3 (181-266) domain previously investigated^28^, the presence of the additional linker residues 267-277 in our D3 construct provokes considerable μs-ms conformational exchanges or/and dynamic association^33,34,39^, which lead to a significant line broadening observed on the current D3 construct (181-277) (Fig. 2B). Thus we proposed that slightly different from the previously-observed dimer formed by the D3 (181-266) (state I of Fig. 2B)^28^, the current D3 domain (181-277) forms a more dynamic dimer due to the presence of the extra linker residues 267-277, which may compete interactions with the dimeric interface of the D3 domain (state II of Fig. 2B).

By contrast, our D5 construct has many sharp up-field 1D peaks and a well-dispersed HSQC spectrum (Fig. 2C), which is very similar to that previously published^28^. DLS results reveal that the D5 domain has an apparent molecular weight of ∼11 kDa at protein concentrations of 100 and 500 μM, suggesting that our D5 domain is also a monomer in solution. Therefore, we decided to carry out detailed NMR structural and dynamic studies on the D5 domain.

### Thermal unfolding of the full-length protein S1

To assess their thermodynamic stability, we conducted thermal unfolding experiments on the S1 protein and its D3 and D5 domains as before on the FUS RRM domain^34^. As the full-length S1 protein contains two free Cys residues, a high-concentration of DTT (2 mM) was added to prevent the formation of non-native disulfide bridge during thermal unfolding. Consequently we were unable to monitor its far-UV CD spectra during thermal unfolding because the noise is unacceptably high in the presence of DTT at 2 mM. Instead, we carried out thermal unfolding by monitoring its near-UV CD and intrinsic UV fluorescence spectra. At a protein concentration of 15 μM in the low salt buffer with 2 mM DTT, upon increasing temperature from 20 to 90 °C, the unfolding occurred as monitored by its near-UV CD (Fig. 3A) and intrinsic UV fluorescence (Fig. 3B) spectra. Interestingly, although no sample precipitation was detected, the unfolding process appears to lack of cooperativity as followed by near-UV CD (Fig. 3C) and intrinsic UV fluorescence (Fig. 3D) probes. Furthermore, the unfolding process is highly irreversible as the sample cooled down to 20 °C after thermal unfolding has the near-UV CD (Fig. 3A) and intrinsic UV fluorescence (Fig. 3B) spectra very different from those of the initial samples. Moreover, in the high salt buffer, the full-length S1 sample at 15 μM became precipitated at ∼60 °C (Fig. 3E), as reported by the overflow of the CD machine HT voltage (> 600), which indicates that the detector is saturated due to the strong scattering light produced by aggregates. At a protein concentration of 50 μM, the full-length S1 sample got precipitated at ∼50 °C in the low salt buffer (Fig. 3F), and at ∼40 °Cin the high salt buffer (Fig. 3G). The results indicate that the protein S1 is more prone to aggregation in the high salt buffer mimicking cellular environment.

**Figure 3.**
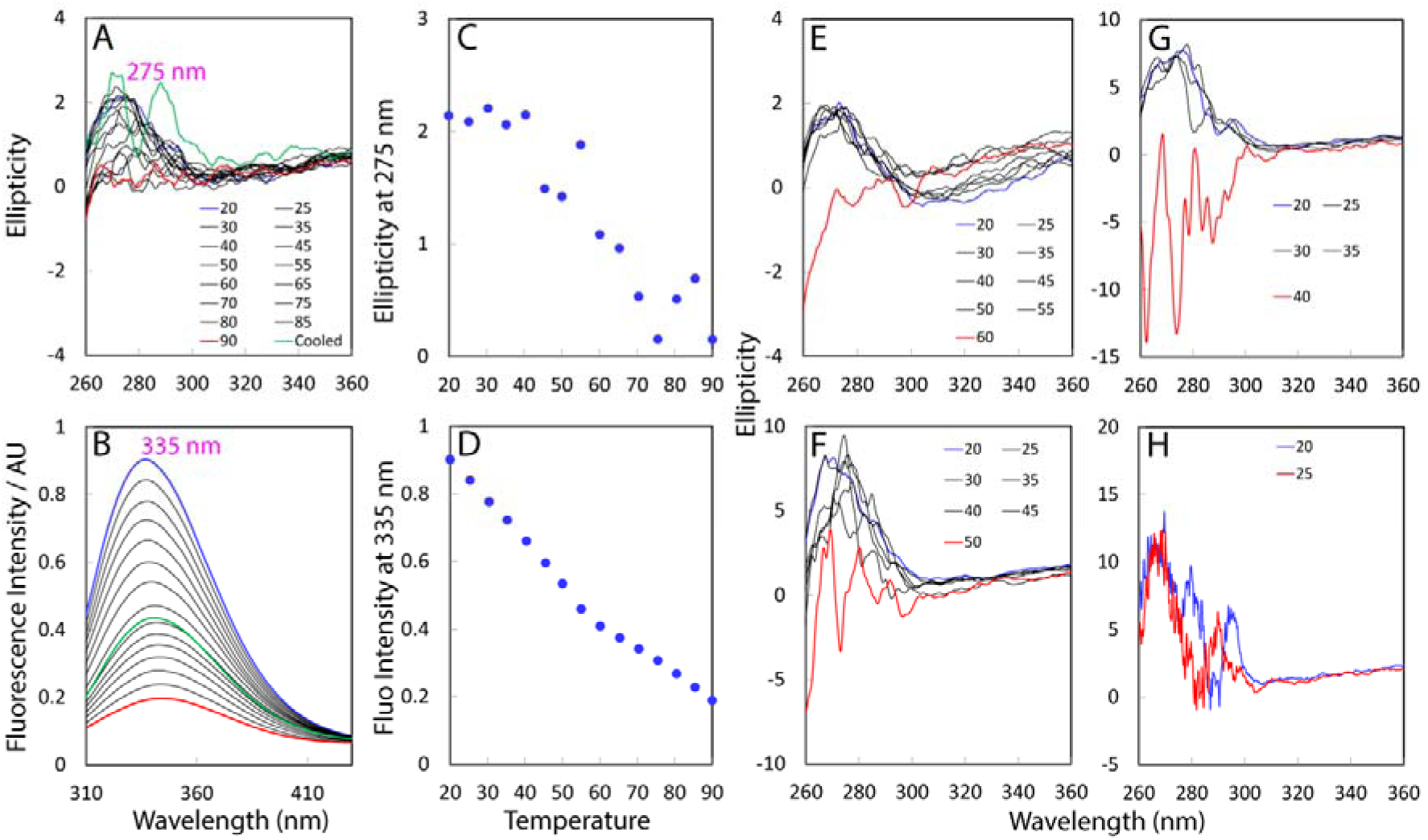
Thermal unfolding of the full-length S1 protein. Near-UV CD (A) and intrinsic UV fluorescence (B) spectra of the full-length S1 protein collected in 10-mm curvet at a concentration of 15 μM in the low salt buffer with temperatures increasing from 20 to 90 °C. The thermal unfolding curves as reported by the changes of ellipiticity at 275 nm (C); and of fluorescence intensity at 335 nm (D). (E) Near-UV CD spectra at a concentration of 15 μM in the high salt buffer. Near-UV CD spectra at a concentration of 50 μM in the low salt buffer (F), and in the high salt buffer (G). (H) Near-UV CD spectra at a concentration of 100 μM in the low salt buffer. The spectra for the samples with significant precipitation detected are colored in red.

We also tested the sample at 100 μM. However, even in the low salt buffer, the machine HT voltage was already larger than 600 at 20 °C and the near-UV spectra have unacceptably high noises (Fig. 3H). Consequently no thermal unfolding of higher protein concentrations was further performed.

### Thermal unfolding of the individual D3 and D5 domains

As both D3 and D5 domains contain no Cys residues, we were able to conduct the thermal unfolding on D3 by monitoring its far-UV CD (Fig. 4A) and fluorescence spectra (Fig. 4B) at a protein concentration of 15 μM in the low salt buffer. Upon increasing temperature, the D3 domain became gradually unfolded without precipitation but again showed no significant cooperativity of unfolding, as clearly indicated by the unfolding curves of both CD ellipticity at 222 nm (Fig. 4C) and fluorescence intensity at 342 nm (Fig. 4D). Noticeably, its far-UV CD spectrum at 90 °C is very different from that of a highly unfolded protein, implying that the secondary structures were not completely denatured, or/and the D3 domain became inter-molecularly associated^34^. Indeed, although the D3 domain has no Cys residue, its CD and fluorescence spectra for the sample cooled down to 20 °C are very different from those of the initial sample, indicative of the irreversibility of the unfolding. The results imply that the D5 domain might also form dynamic oligomers upon being unfolded and consequently failed to refold to the initial structure upon being cooled down, exactly as we previously observed on the FUS RRM domain^34^. At a protein concentration of 50 μM, the D3 domain sample became precipitated at ∼70 °C in the low salt buffer (Fig. 4E) and at ∼65 °C in the high salt buffer (Fig. 4F).

**Figure 4.**
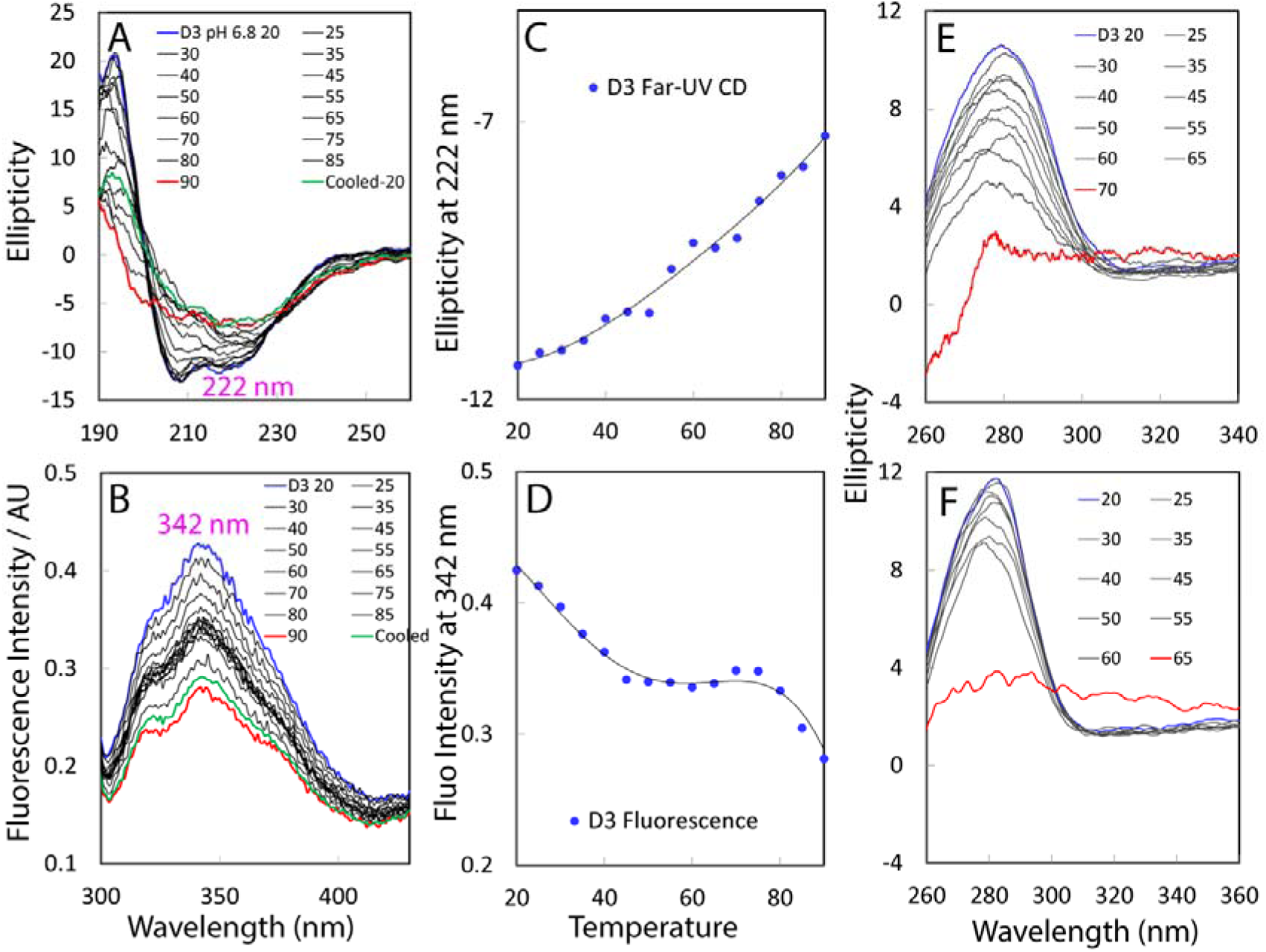
Thermal unfolding of the D3 domain. Far-UV CD (A) and intrinsic UV fluorescence (B) spectra of the D3 domain collected in 1-mm curvet at a concentration of 15 μM in the low salt buffer. The thermal unfolding curves as reported by the changes of ellipiticity at 222 nm (C); and of fluorescence intensity at 342 nm (D). Near-UV CD spectra of the D3 domain at a protein concentration of 50 μM collected in 10-mm curvet at 25 °C in the low salt buffer (E); and the high salt buffer (F). The spectra for the samples with significant precipitation are colored in red.

We also conducted the same unfolding experiments on the D5 domain. At 15 μM, the D5 domain also became gradually unfolded (Fig. 5A and 5B) without significant cooperativity of unfolding (Fig. 5C and 5D). Again, its far-UV CD spectrum at 90 °C is very different from that of a highly unfolded protein. Like the D3 domain, the unfolding of the D5 domain is also highly irreversible. At 50 μM, the D5 domain sample became precipitated at ∼60 °C in the low salt buffer (Fig. 5E), and at ∼55 °C in the high salt buffer (Fig. 5F).

**Figure 5.**
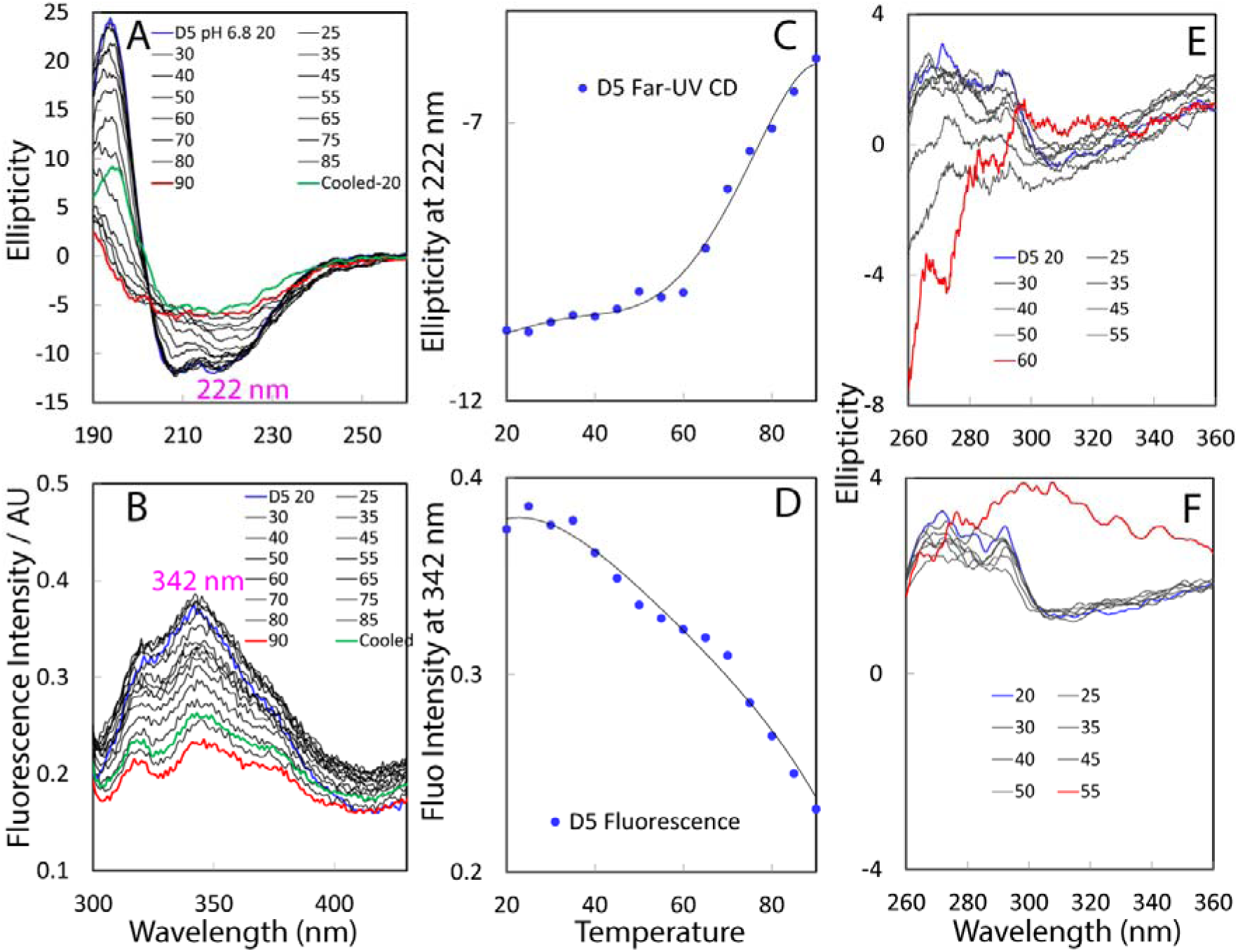
Thermal unfolding of the D5 domain. Far-UV CD (A) and intrinsic UV fluorescence (B) spectra of the D5 domain collected in 1-mm curvet at a concentration of 15 μM in the low salt buffer. The thermal unfolding curves as reported by the changes of ellipiticity at 222 nm (C); and of fluorescence intensity at 342 nm (D). Near-UV CD spectra of the D5 domain at a protein concentration of 50 μM collected in 10-mm curvet at 25 °C in the low salt buffer (E); and in the high salt buffer (F). The spectra for the samples with significant precipitation are colored in red.

### Backbone dynamics on different time scales

Our D3 domain with the extra linker had very broad NMR resonance signals and many HSQC peaks were undetectable (Fig. 2B), thus preventing further high-resolution NMR studies. Here we focused on determining the NMR structure and dynamic of the D5 domain.

By analyzing a pair of triple resonance NMR spectra HN(CO)CACB/CBCA(CO)NH, as well as ^15^N-edited HSQC-TOCSY/ HSQC-NOESY, we have accomplished the sequential assignment of the D5 (350-443) domain except for the NH and ^15^N chemical shifts of Pro353 and Pro440. Fig. S2A presents the (ΔCα-ΔCβ) chemical shifts, which represent a sensitive indicator of the secondary structures of both folded and disordered proteins^40^. Many residues have very large (ΔCα–ΔCβ), characteristic of a well-folded protein, suggesting that the D5 domain is well-folded. On the other hand, analysis of NOESY spectra revealed that unlike other four S1 motifs whose NMR structures could be determined with a large number of long-range NOEs (Fig. S1)^29,30^, although there exist characteristic NOEs defining secondary structures (Fig. S2B), only 55 long-range NOEs (Table S1) could be identified over a small portion of residues (Fig. S2C). This number is much less than what we previously obtained from other well-folded proteins, as exemplified by a 37-residue charybdotoxin with an α/β motif stabilized by three disulfide bridges and its two-disulfide derivative^41,42^, an 81-residue viral protein VP9 with an α/β ferredoxin fold without any disulfide bridge^43^, as well as an 183-residue ectodomain of EphA4 receptor with a β-barrel constrained by two disulfide bridges^44^. This is reminiscent of the ubiquitin-like folds of the TDP-43 N-terminal domain^39^ and FAT10^45^, which were also short of long-range NOEs. Strikingly, both ubiquitin-like folds have been characterized to undergo significant conformational dynamics which accounted for their intrinsic proneness to aggregation^39,45^.

So we decided to assess its conformational dynamics on different time scales. We first collected ^15^N backbone relaxation data T1, T2 and {^1^H}-^15^N steady-state NOE (hNOE). However, likely due to conformational exchanges, the HSQC peaks for measuring T1 and T2 were highly broad at some delay times as we previously found^39,45^, as such we were unable to obtain T1 and T2 values. Fig. 6A presents its residue-specific hNOE which provides a measure to the backbone flexibility on the ps-ns time scale^33,34,39,40,44^. The D5 domain has positive hNOE values for all residues except for Lys350 with an average value of 0.78, indicating that the D5 domain has significantly-restricted backbone motions on the ps-ns time scale.

**Figure 6.**
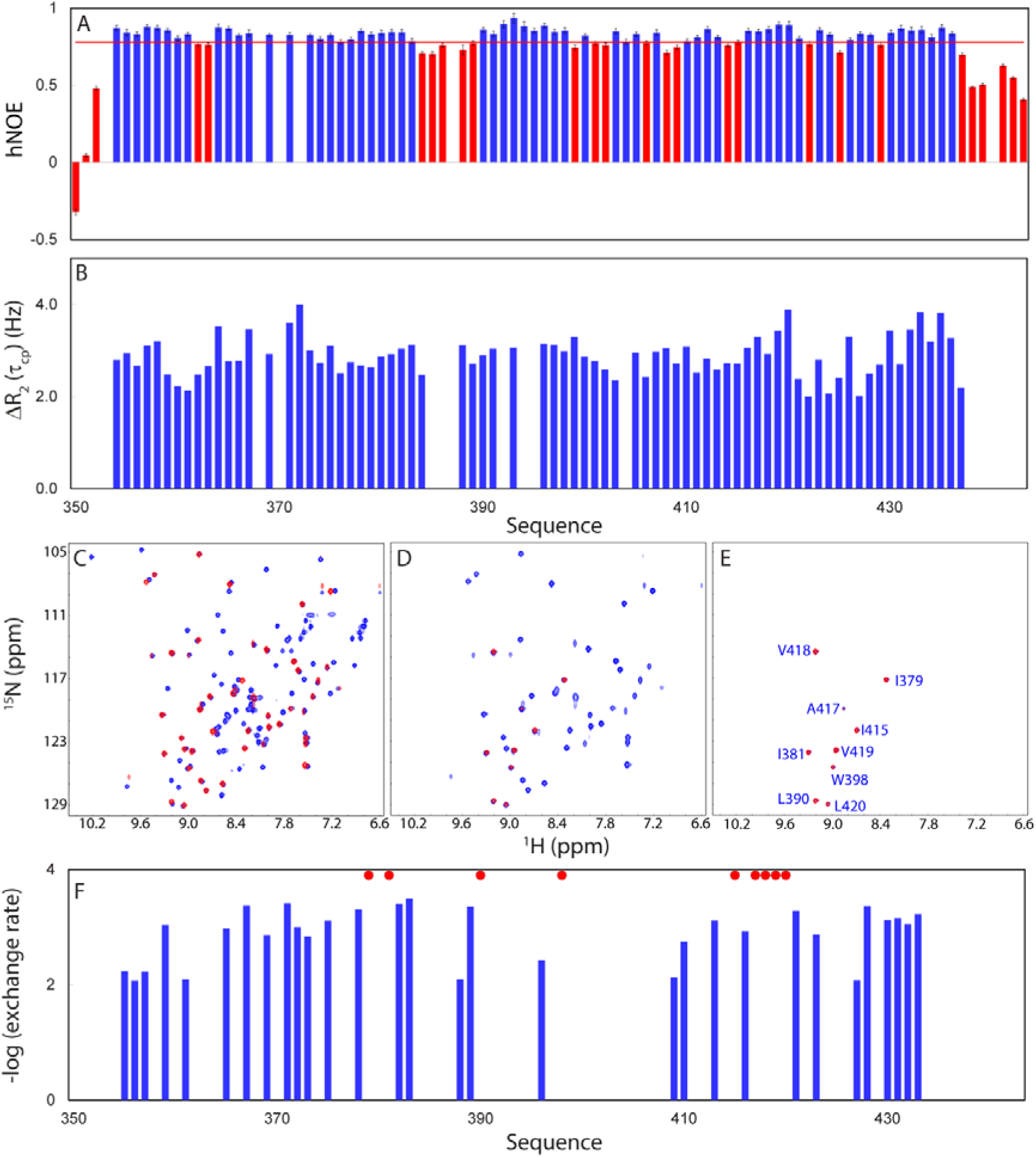
NMR backbone dynamics of the D5 domain on different time scales. (A) {^1^H}-^15^N heteronuclear steady-state NOE (hNOE) values with the average value of 0.78 displayed as the red line. The bars with values < the average are colored in red. (B) CPMG dispersion data as represented by the differences of effective transverse relaxation rate R_2_(t_cp_) at 40 and 960 Hz respectively. Only residues with ΔR_2_(t_cp_) > 2 Hz are displayed. (C) Superimposition of HSQC spectra of the D5 domain acquired at 25 °C at a protein concentration of 200 μM in the high salt buffer (blue); and 10 minutes after re-dissolving the lyophilized sample into D_2_O (red). (D) Superimposition of HSQC spectra of the D3 domain of the same sample acquired 10 minutes (blue) and 4 hours after re-dissolving the lyophilized sample into D_2_O (red). (E) Superimposition of HSQC spectra of the D3 domain of the same sample acquired 4 hours (blue) and 24 hours (red) after re-dissolving the lyophilized sample into D_2_O (red). (F) Logarithmic plot of the experimentally measured hydrogen-deuterium exchange rates of the D5 domain. The red circles are used to indicate nine slow-exchange-rate residues.

We next obtained its CPMG-based relaxation dispersion data (Fig. 6B), and interestingly the majority of the D5 residues have ΔR_2_(τ_cp_) > 2.0 Hz, thus suggesting the existence of the μs-ms conformational exchanges over the whole molecule. However, due to the relatively small values (all < 4 Hz), we were unable to quantitatively fit the data to obtain exchange parameters as we previously performed on the EphA4 receptor in the free state and in complex with two different small molecules^44^.

We further carried out NMR hydrogen/deuterium (H/D) exchange experiments to assess the backbone dynamics on the min-hr time scale. It is well-established that in solution the labile hydrogens such as amide protons of proteins are continually exchanging with the solvent at different rates, depending on a variety of factors including their exposure to the solvent or/and their involvement in H-bonds. Consequently, amide H/D exchange results offer a sensitive reflection of the exposure degree of amide protons to the bulk solvent^46–51^. As seen in Fig. 6C, upon subjecting to H/D exchange, all side-chain amide protons of Asn or Gln, as well as ∼50% of backbone amide protons of the D5 residues have completely exchanged with deuterium within the dead time of the first HSQC experiment (10 min). After 4 hr, more amide protons exchanged and consequently only 9 residues accounting for ∼9.6% of the total residues have persisted HSQC peaks (Fig. 6D), thus categorized to be slow-exchange-rate residues^46–51^. After 24 hr, those nine HSQC peaks still persisted although their intensity further reduced (Fig. 6E).

With the residue-specific H-D exchange data at different time points, we attempted to fit out the exchange rates for the residues excepted for those with the complete H-D exchange within the dead time as well as those remaining after 24 hr (Fig. 6F). Strikingly, except for the nine slow-exchange-rate residues, all D5 residues have –log(exchange rate) values < 4, much less than those of the residues in the secondary structure regions of the sperm whale myoglobin which range from 6 to 8, only close to those of its loop/linker residues whose values are ∼4^47^.

We also conducted the H-D exchange experiments on the D3 domain despite having no sequential assignments due to the significant broadening of NMR peaks (Fig. 3B). Similar to the D5 domain, with the dead time, ∼60% of HSQC peaks completely disappeared (Fig. 7A). After 1 hr, more peaks disappeared (Fig. 7B), while after 4 hr, only a total of 17 HSQC peaks could be detected (Fig. 7C). After 24 hr, only 14 HSQC peaks retained. Noticeably, out of the remaining 14 HSQC peaks, at least six could be assigned to the side-chain amide protons of Asn or Gln residues (Fig. 7D). This is unexpected as for most cases, the side-chain amide protons of Asn or Gln residues are highly accessible to the bulk solvent and consequently they completely exchanged with deuterium even within the dead time, as evident from the result of the D5 domain (Fig. 6C). This observation implies that those side-chain amide protons of the D3 Asn or Gln residues are highly buried or/and involved in the hydrogen bonding, probably due to their involvement in the dimerization observed previously^28^ and here.

**Figure 7.**
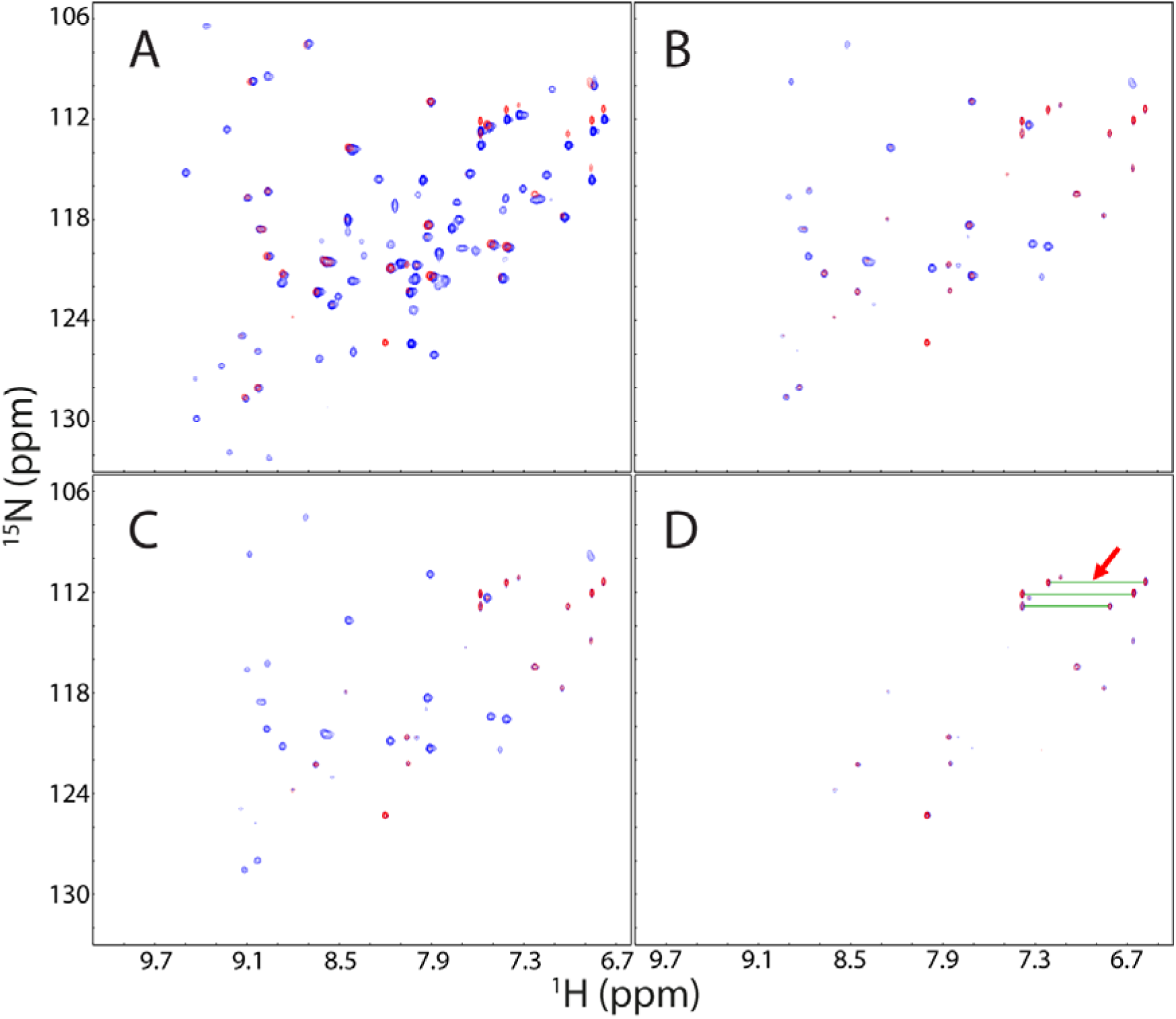
NMR hydrogen-deuterium (H-D) exchange experiments of the D3 domain. (A) Superimposition of HSQC spectra of the D3 domain acquired at 25 °C at a protein concentration of 200 μM in the high salt buffer (blue) and 10 minutes after re-dissolving the lyophilized sample into D_2_O (red). (B) Superimposition of HSQC spectra of the D3 domain of the same sample acquired 10 minutes (blue) and 1 hours after re-dissolving the lyophilized sample into D_2_O (red). (C) Superimposition of HSQC spectra of the D3 domain of the same sample acquired 10 minutes (blue) and 4 hours re-dissolving the lyophilized sample into D_2_O (red). (D) Superimposition of HSQC spectra of the D3 domain of the same sample acquired 4 hours (blue) and 24 hours (red) after re-dissolving the lyophilized sample into D_2_O (red). The red arrow is used to indicate the side-chain amide protons of Gln or Asn residues which remain un-exchanged even 24 hr after re-dissolving the lyophilized sample into D_2_O.

Although all six S1 repeats of *E. coli* ribosomal protein S1 assume the conserved S1 motif with the β-barrel fold, the D3 and D5 domains appear to have higher min-hr dynamics than other four S1 repeats, whose NMR structures have been previously determined with the input of many hydrogen-bond constraints derived from the slow-exchange-rate backbone amide protons^29,30^. Furthermore, the D3 and D5 domains have the min-hr dynamics not only much higher than those of the sperm whale myoglobin with the paradigm globin fold of eight helices^47^, but also higher than those of the viral protein VP9 with an α/β ferredoxin fold^43^, as well as the ephrin-B2 ectodomain^48^, MSP domain of VAPB protein^49^, EphA4 and EphA5 ectodomains^50,51^, which all adopt the β-barrel folds.

### NMR structures of the D5 domain

We calculated 50 NMR structures of the D5 domain by CYANA software package^52^ with NOE-derived distance and TALOS-based dihedral angle^53^ restraints as shown in Table S1. Subsequently the five structures with the lowest target functions were selected for further refinement with AMBER force field^54^. The five refined structures are shown in Fig. 8A and the lowest energy one in Fig. 8B. The D5 domain adopts the conserved S1 fold with its five β-strands formed respectively over residues Gly364-Gly369, Ile379-Gly382, Asp388-Val391, Asp413-Asp423, and Glu427-Leu431 (Fig. 8B). Interestingly, the D5 domain also contains additional four helices over residues Trp354-His361, His392-Ile396, Gly402-Arg407 and Val433-Leu436 respectively. However, the numbers, locations and lengths of helices appear to be highly variable in the five S1 repeats (D1, D2, D4, D5 and D6) with NMR structures solved, and in particular the D4 domain (PDB code of 2KHI) even contains no helix at all (Fig. S1)^29^. For the D5 domain, while its β-barrel regions are well-defined with the backbone RMSD value of 0.54 Å (Table S1), the other regions show large variations in different structures (Fig. 8A).

**Figure 8.**
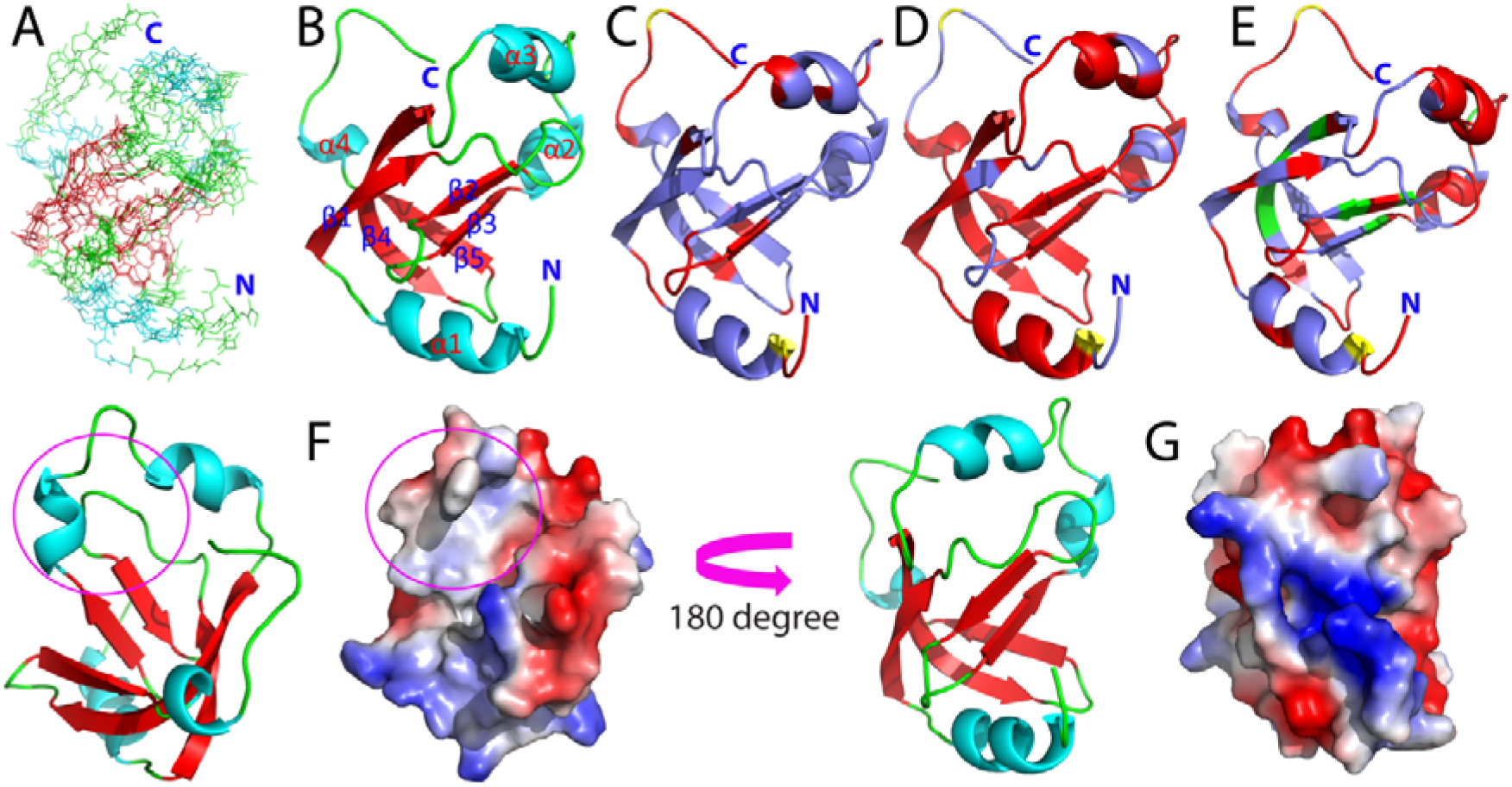
Visualization of NMR structures and dynamics of the D5 domain. (A) Superimposition of the five selected structures. Red: β-barrel regions; cyan: helices; and green: loops. (B) The lowest energy structure with the secondary structure elements labelled. (C) NMR structure with the residue-specific hNOE mapped back, yellow: Pro residues; red: residues with hNOE < the average (0.78). (D) NMR structure with the residue-specific CPMG data mapped back, yellow: Pro residues; red: residues with ΔR_2_(t_cp_) > 2 Hz. (E) NMR structure with the residue-specific H-D exchange data mapped back, yellow: Pro residues; red: residues with their backbone amide protons completely exchanged within the dead time; green: residues with their HSQC peaks persisted four hours after re-dissolving the lyophilized D5 sample into D_2_O; and light blue: residues with their amide protons exchanged between the dead time and four hours. (F) Ribbon structure and electrostatic potential surface of the D5 domain, and with their orientations rotated 180° along the z axis (G).

Upon mapping the residue-specific hNOE (Fig. 6A) back to the D5 structure, it appears that in addition to the N-/C-terminal and loop residues, some residues in β-strands (β3, β4 and β5) and α-helices (α3 and α4) also have hNOE values less than the average (0.78), implying that these residues also have relatively high conformational dynamics on the ps-ns time scale (Fig. 8C). Furthermore, mapping of the CPMG data (Fig. 6B) back to the structure reveals that except for the N-/C-terminal and some residues with high ps-ns conformational dynamics, most D5 residues have conformational exchanges on the μs-ms time scale (Fig. 8D). Fig. 8E shows the D5 structure with the H-D exchange results mapped back. Strikingly, the residues with their backbone amide protons completely exchanged with the dead time are not just limited to the terminal and loop residues, but located over the whole molecule which include all five β-strands and four α-helices (Fig. 8E). Furthermore, out of nine slow-exchange-rate residues, two are located within the second β-strand, one within the third β-strand while five within the forth β-strand. Interestingly, one is located in the loop connecting the second and third helices, but none within the helical region (Fig. 8E). The results clearly indicate that in addition to terminal and loop residues, most residues in the secondary structure regions are also highly accessible to the bulk solvent. As a result, even the conserved β-barrel of the D5 domain has significant dynamics on the min-hr time scale (Fig. 8E). This is most likely resulting from the “global breathing” motions via “unfolding equilibria”^46–51^, which has been previously found to exist to different degrees on other proteins with the β-barrel folds, including the ephrin-B2 ectodomain^48^, MSP domain of VAPB protein^49^, EphA4 and EphA5 ectodomains^50,51^.

It is considerably interesting to note that the D5 domain has many hydrophobic patches (Fig. 8F and 8G). In particular, a large hydrophobic pocket is observed mainly over the surface constituted by the second/third helices and loops (Fig. 8F). Moreover, given the existence of globular μs-ms dynamics and particularly high min-hr “global breathing” motions, the structurally-buried hydrophobic patches/cores in the D5 domain should be at least dynamically accessible to drive the irreversible inter-molecular association/aggregation under some circumstances.

## Discussion

It had been widely thought that reproduction by asymmetric division is a prerequisite for aging^55^ until the recent discovery that even *E. coli* cells progressively decline in growth rate and reproductive ability^16,17^, thus revealing that no life can escape the attack of aging. One universal characteristic associated with aging of organisms from human being down to *E. coli* cells is aggregation of hundreds and thousands of non-specific proteins^12–23,56–58^. However, due to the extreme complexity, it remains challenging to address whether the aggregated proteins trigger aging of mammalian organisms through “gain of toxic functions” by which aggregation-prone proteins such as α-synuclein, TDP-43 and FUS initiate neurodegenerative diseases^4,12,33,34,39,59^. By contrast, it appears that protein aggregates do impose damaging effects to *E. coli* cells because obviously different from the daughter cell free of protein aggregates, the mother cell holding the aggregates has a reduced growth rate, decreased offspring production, and an increased incidence of death^16,17^. Most beneficially, as compared with mammalian aging for which nine tentative hallmarks have been enumerated^56^ and more than 300 different theories have been proposed^60^, so far the most significant biomarker for cellular aging and rejuvenation of *E. coli* cells appears to be only protein aggregation. Consequently, *E. coli* offers a simple but an excellent model for addressing the fundamental relationship between protein aggregation and cellular aging, which includes at least two key questions: 1) what are the mechanisms by which so many proteins become aggregated during aging; and 2) whether aggregated proteins gain toxic functions responsible for aging of *E. coli* cells.

In the present study, we attempted to explore the mechanisms for aggregation of *E. coli* proteins without any covalent modifications of sequences such as side-chain modifications and fragmentation triggered by over-oxidation^57^, through a detailed biophysical characterization of conformations, unfolding properties and dynamics of ribosomal protein S1 and its isolated D3 and D5 domains. We are particularly interested in understanding the observed paradox that while *E. coli* ribosomal protein S1 was characterized to be well-folded and highly soluble, it has been also found in all lists of aggregated proteins of *E. coli* cells, even in the healthy cells without significant oxidative stress^23^. The obtained results reveal that the aggregation of the protein S1 is mediated by dynamics at both inter-domain interaction and individual domain levels.

### The roles of dynamics of inter-domain interactions in aggregation

Firstly, ANS-binding results indicate that the full-length S1 and its D3/D5 domains contain significantly exposed hydrophobic patches comparable to those characteristic of the molten globule state, an aggregation-prone intermediate of protein folding, which, unlike the native state of a well-folded protein with hydrophobic cores tightly buried, still has its hydrophobic cores loosely packed, thus accessible to binding ANS^35,36,61–63^. Our NMR characterization also confirms the recent reports that the well-folded six S1 repeats have dynamic inter-domain interactions^27,28^. These two characteristics of the protein S1 appear to be the structural basis essential for the optimized implementation of its biological functions, but on the other hand unavoidably results in its proneness to aggregation under some circumstances. Briefly, the functional roles of the protein S1 require its six S1 repeats rather individually available for differential binding to a variety of partners such as ribosome and nucleic acids. However, as some of the six repeats (at least D3 and D5 as shown here) contain significantly exposed hydrophobic patches, adopting a highly extended conformation with all six S1 repeats completely exposed to the bulk solvent would unavoidably lead to severe aggregation in particular in cellular environments with high salt concentrations. As a trade-off, the protein S1 establishes dynamic inter-domain interactions, which on the one hand can shield the hydrophobic patches from being completely accessible to the bulk solvent in the free state, but on other hand allow a rapid structural rearrangement to release the dynamically-packed S1 repeats for binding upon availability of the partners. This strategy appears reducing but unable to completely eliminate the proneness of S1 to aggregation. This is reminiscent of the ALS-causing TDP-43 intrinsically prone to aggregation, which has “open” and “closed” conformational states mainly mediated by dynamic inter-domain interactions via hydrophobic regions^64^ and its disruption by ALS-causing cleavage leads to a significantly enhancement of aggregation^65^.

The dynamic nature of the inter-domain interactions rationalizes the experimental results that the protein S1 was significantly prone to aggregation at slightly high temperatures and in the high salt buffer. The thermal unfolding results show that in the high salt buffer mimicking cellular environment, the protein S1 became precipitated even at 40 °C at a protein concentration of 50 μM comparable to the ribosome concentration in an *E. coli* cell^26^ (Fig. 3G). Noticeably, this temperature is very close to the physiological temperature for *E. coli* optimized growth and still much lower than those for the individual domains such as D3 and D5 to be significantly unfolded. Nevertheless, this temperature appears sufficient to largely disrupt the dynamic interactions among the S1 repeats and consequently increase the probability to expose hydrophobic patches for driving inter-molecular clustering/aggregation, particularly in high salt buffer. In 2005, we found that even previously-thought “completely insoluble” proteins could in fact be solubilized in the salt-minimized water^10–12,66^. Further analysis decoded that unlike the well-folded proteins following the “Salting-in” rule established a century ago that protein solubility increases upon adding salts over the range of low salt concentrations (usually <300–500 mM)^67^, for the “completely insoluble” proteins with significant exposure of hydrophobic patches, a very low salt concentration is sufficient to trigger aggregation/insolubility due to screening out of the repulsive electrostatic interaction by non-specific ionic strength, or/and specific anion-binding to proteins^10–12,68^.

Remarkably, very recently it was deciphered that the oligomerization of the aggregation-prone but not “completely insoluble” prion protein was also predominantly mediated by salt concentrations through both non-specific ionic strength or/and specific anion-binding to proteins^38^. Therefore, the salt-enhanced aggregation of *E. coli* ribosomal protein S1 is expected to follow the same mechanism previously established for “completely insoluble” and aggregation-prone protein10-12,33,34,39,45,49,64-66,68.

### The roles of dynamics of individual domains in aggregation

Secondly, high-resolution NMR characterizations decipher the structural and dynamic basis for the D3 and D5 domains to have significantly exposed hydrophobic patches. For example, despite adopting the conserved β-barrel fold with significantly-restricted backbone motions on the ps-ns time scale, the D5 domain has global μs-ms conformational dynamics and extremely high min-hr “global breathing” motions. As a consequence, the high dynamics particularly on the μs-ms and min-hr time scales could even render the structurally-buried hydrophobic patches/cores to become dynamically accessible, thus explaining the observation that the well-folded D5 domain has an unexpectedly high ANS binding. Furthermore, the existence of many structurally- or/and dynamically-exposed hydrophobic patches/cores may also account for the unfolding irreversibility of the D3 and D5 domain because these hydrophobic patches may drive irreversible inter-molecular associations even at the early stage of unfolding, as we previously observed on the RRM domain of another ALS-causing protein FUS which is also intrinsically aggregation-prone^34^. While it remains to be explored whether the observed high dynamics of the D3 and D5 domains are required for their biological functions, the high dynamics at the individual domain level certainly contribute to the proneness of S1 to aggregation.

Noticeably, it appears that compared to the all-helical proteins such as myoglobin^47^, the proteins adopting β-rich architectures have relatively high conformational dynamics most likely because both secondary and tertiary structures of β-rich proteins are highly dependent of the stabilization from the long-range interactions, which are more vulnerable to various perturbations than that from the local interactions stabilizing the helical structures^36,61,69,70^. Indeed, even we have previously identified the presence of significant conformational dynamics on differential time scales associated with the proteins adopting β-rich architectures, which include FUS RRM domain^34^, the ubiquitin-like folds of TDP-43^39^ and FAT10^45^, and MSP domain of VAPB protein^49^, as well as the ephrin-B2^48^, EphA4^50^ and EphA5^51^ ectodomains, which all adopt the β-barrel folds even stabilized by two disulfide bridges. These together imply that like protein S1, due to the existence of relatively high dynamics, a large portion of β-rich proteins could be also prone to aggregation *in vivo* with high salt concentrations even without any covalent modifications. This might at least partly account for the observation that the aggregated proteins during aging are over-represented by proteins with β-rich architectures^12–15^.

## Conclusion

Our biophysical studies here reveal that high dynamics of *E. coli* protein S1 at both inter-domain interaction and individual domain levels lead to significant exposure of hydrophobic patches/cores, which drive aggregation of the protein S1 in cell with high salt concentrations. This thus rationalizes the paradoxical phenomenon that while the protein S1 is well-folded and very soluble *in vitro* in relatively low salt buffers, it could be prone to aggregation in cell even without any covalent modifications. Furthermore, upon over-oxidation associated with aging to trigger fragmentation, some fragments of the protein S1 became “completely insoluble” in buffers^71^.

As proteins from prokaryotic and eukaryotic organisms have no fundamental difference and most of them, if not all, have conformational dynamics on certain time scales, the mechanism we decoded here for the protein S1 by which dynamics critically mediate protein aggregation most likely also operates in other proteins of *E. coli*, as well as other organisms including human being. However, in eukaryotic organisms, the physiologically relevant post-translational modifications such as phosphorylation may also mediate protein aggregation by changing protein structures and dynamics of individual domain or/and inter-domain interactions^64,65,72^.

## Methods

### Preparation of recombinant proteins

The DNA fragments encoding the full-length *E. coli* ribosomal S1 protein was amplified by PCR reactions from *E. coli* BL21 cells (Novagen) and subsequently its two dissected domains D3 (181-277) and D5 (350-443) were amplified by PCR reactions from the full-length S1 protein gene. The DNA fragments were cloned into a modified vector pET28a we previously used for the 526-residue FUS and its dissected domains^34^. The expression vectors were transformed into and overexpressed in *E. coli* BL21 (DE3) cells (Novagen). The recombinant proteins of the full-length S1 protein and its D3 domain was highly soluble in supernatant, while the D5 domain was found in inclusion body if induced by 1 mM IPTG at 37 °C, exactly as previously reported^28^. However, if we performed cell culture to express the D5 domain at 20 °C with induction by 0.2 mM IPTG, a soluble portion was found in supernatant. Consequently, in the present study all three recombinant proteins have been successfully purified by Ni^2+^-affinity column (Novagen) under native condition, followed by a further purification by FPLC on gel-filtration columns.

The generation of the isotope-labelled proteins for NMR studies followed a similar procedure except that the bacteria were grown in M9 medium with the addition of (^15^NH_4_)_2_SO_4_ for ^15^N labeling and (^15^NH_4_)_2_SO_4_ /[^13^C]-glucose for double labelling. The purity of the recombinant proteins was checked by SDS-PAGE gels and their molecular weights were verified by a Voyager STR matrix-assisted laser desorption ionization time-of-flight-mass spectrometer (Applied Biosystems) as we previously performed^33,34,44,45,48–51^. The concentration of protein samples was determined by the UV spectroscopic method in the presence of 8 M urea. Briefly, under the denaturing condition, the extinct coefficient at 280 nm of a protein can be calculated by adding up the contribution of Trp, Tyr and Cys residues^73^.

### CD and fluorescence experiments

All circular dichroism (CD) experiments were performed on a Jasco J-1500 spectropolarimeter equipped with a thermal controller. Far-UV CD spectra were collected in 1-mm curvet at a protein concentration of 15 μM in the low salt buffer (1 mM phosphate buffers at pH 6.8), while near-UV CD spectra were acquired in 10-mm curvet in either low salt or high salt (50 mM phosphate buffers at pH 6.8 with 200 mM NaCl) buffers, because much high protein concentrations are needed to obtain high-quality near-UV spectra. Data from five independent scans were added and averaged. Due to the presence of two Cys residues, for the samples of the full-length protein S1, DTT was further added to reach a final concentration of 0.5 mM to prevent the formation of non-native disulfide bridges.

Thermal unfolding of S1 and its D3 and D5 domains was carried out by increasing temperature from 20 to 90 °C at several protein concentrations and in either low salt or high salt buffers on a Jasco J-1500 spectropolarimeter, which is able to simultaneously acquire both CD and fluorescence spectra during the thermal unfolding. For the samples of the full-length protein S1, DTT was added to reach a final concentration of 2 mM. Intrinsic UV fluorescence spectra were collected between 300 and 430 nm with the excitation at 280 nm.

### Dynamic light scattering (DLS) experiments

The apparent molecular weights of the full-length S1, D3 and D5 domains were assessed by use of dynamic light scattering (DLS) at 25 °C on a DynaPro-E-50-830 instrument (Protein Solutions Inc., Lakewood, NJ, USA) as previously conducted^74^. The samples were centrifuged at 15,000 rpm for half an hour to remove tiny particles and air bubbles. For each protein, its apparent relative molecular mass was measured at two protein concentrations: 100 and 500 μM. The values of apparent relative molecular mass were calculated and averaged from at least 10 measurements using the built-in analysis software as we previously conducted^74^.

### ANS Fluorescence

Aliquots of the full-length S1 or D3 or D5 protein solution were taken and added into assay solution to give a final protein concentration of 20 μM in 80 μM ANS in high salt buffer with 2 mM DTT. An Agilent Technologies Fluorimeter was used to record fluorescence spectra between 410 and 700 nm with the excitation at 350 nm.

### NMR experiments

All NMR experiments were acquired on an 800 MHz Bruker Avance spectrometer equipped with pulse field gradient units as described previously^33,34,44,45,48–51^. NMR samples for collecting 1D and HQSC spectra were prepared at a protein concentration of 50 μM in high salt buffer. For NMR samples of the full-length protein, DTT was further added to reach a final concentration of 20 mM to prevent the formation of non-native disulfide bridges as previously used^28^.

For determining residue-specific conformation of the D5 domain, a pair of triple-resonance experiments HNCACB, CBCA(CO)NH were collected for the sequential assignment on a ^15^N-/^13^C-double labelled sample at 300 μM in high salt buffer, which was previously used to mimic cellular conditions for NMR characterization of the D3 and D5 domains^28^. For assigning and collecting NOE connectivities, ^15^N-edited HSQC-TOCSY and HSQC-NOESY experiments were acquired on a ^15^N-labeled sample at 500 μM. NMR data were processed with NMRPipe^75^ and analyzed with NMRView^76^.

### Protein dynamics on different time scales as studied by NMR spectroscopy

^15^N backbone T1 and T1ρ relaxation times and {^1^H}-^15^N steady state NOE intensities were collected on the ^15^N -labeled D5 domain at 25 °C at a concentration of 500 μM on an Avance 800 MHz Bruker spectrometer with both an actively shielded cryoprobe and pulse field gradient units as previously described^33,34,39,50^. {^1^H}-^15^N steady-state NOEs were obtained by recording spectra with and without ^1^H presaturation, a duration of 3 s and a relaxation delay of 6 s at 800 MHz.

^15^N transverse relaxation dispersion experiments were acquired on the ^15^N-labeled D5 domain on a Bruker Avance 800 spectrometer as previously described^33,34,44,50^. A constant time delay (*T*_CP_ = 50 ms) was used with a series of CPMG frequencies, ranging from 40 Hz, 80 Hz, 120 Hz (x2), 160 Hz, 200 Hz, 240 Hz, 320 Hz, 400 Hz, 480 Hz, 560 Hz, 640 Hz, 720 Hz, 800 Hz, and 960 Hz (x2 indicates repetition). A reference spectrum without the CPMG block was acquired to calculate the effective transverse relaxation rate by the following equation:

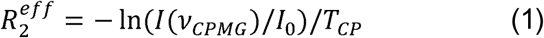

Where I(υ_CPMG_) is the peak intensity on the difference CPMG frequency, I_0_ is the peak intensity in the reference spectra.

Hydrogen-deuterium (H-D) exchange experiments were conducted on both D3 and D5 to gain an initial insight into its dynamic behavior on the min-hr time scale^46–51^. Briefly, an HSQC spectrum was acquired for the ^15^N-labeled D3 or D5 at a protein concentration of 200 μM in high salt buffer and subsequently the sample was lyophilized. The H-D exchange experiments were initiated by re-dissolving the lyophilized sample into D_2_O. The progress of the exchange process between amide protons and deuterium was followed by collecting a series of successive HSQC spectra starting immediately after re-solubilization of the sample in D_2_O. The first HSQC spectrum was collected at 10 min after re-solubilization, and the last spectrum was acquired after 24 hours. H-D exchange progress of individual residues was followed by peak intensities in the HSQC spectra acquired at different time points, and data were fitted by a nonlinear least-squares algorithm to a single exponential decay to obtain residue-specific exchange rates^46,47^.

### Determination of NMR structures of the D5 domain

NMR structures of the D5 domain were determined as we previously carried out on other proteins^43–45^. Briefly, 50 NMR structures were calculated by CYANA^52^ with distances derived from NOE connectivities and dihedral angel restraints by TALOS^53^. The five lowest target-function structures with no distance violation > 0.5 Å and dihedral angel violation > 5° were selected for further refinement in Amber99sb-ildn force field and GROMACS version 4.5.3 was used to perform the energy minimization^54^. The NMR structures of the D5 domain and associated NMR data were deposited respectively in PDB (ID of 5XQ5) and in BMRB (ID of 36095). The quality of the refined structures was checked by PROCHECK-NMR^77^ and displayed by PyMOL Molecular Graphics System, version 0.99rc6 (Schrödinger, LLC).

## Acknowledgements

This study is supported by Ministry of Education of Singapore (MOE) Tier 2 grant MOE2015-T2-1-111 to Jianxing Song. The funders had no role in study design, data collection and analysis, decision to publish, or preparation of the manuscript.

## Author Contributions

Conceived and designed the experiments: JXS; Performed the experiments: YML LZL; Analyzed the data: YML LZL JXS. Prepared figures and wrote the paper: JXS.

## Accession codes

The coordinates of the D5 solution structures and associated NMR data were deposited in PDB (ID of 5XQ5) and in BMRB (ID of 36095) respectively.

## Supplementary information

**Figure S1.**
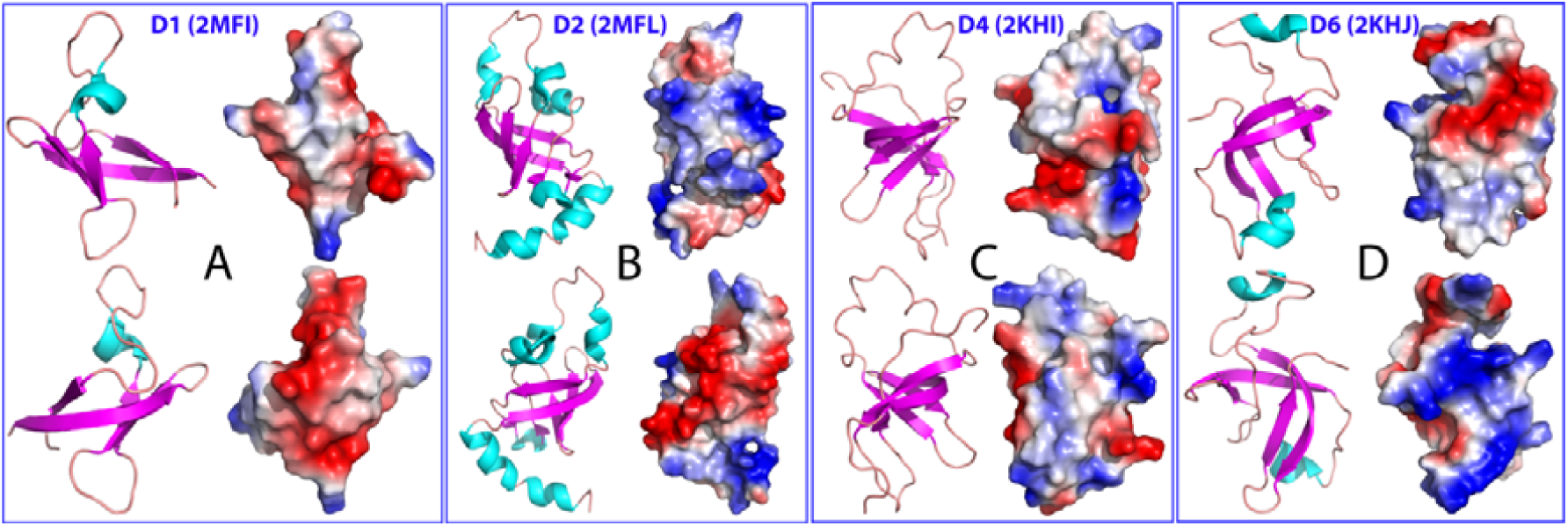
Ribbon structures and electrostatic potential surfaces of four S1 repeats of *E. coli* ribosomal protein S1 previously reported.

**Figure S2.**
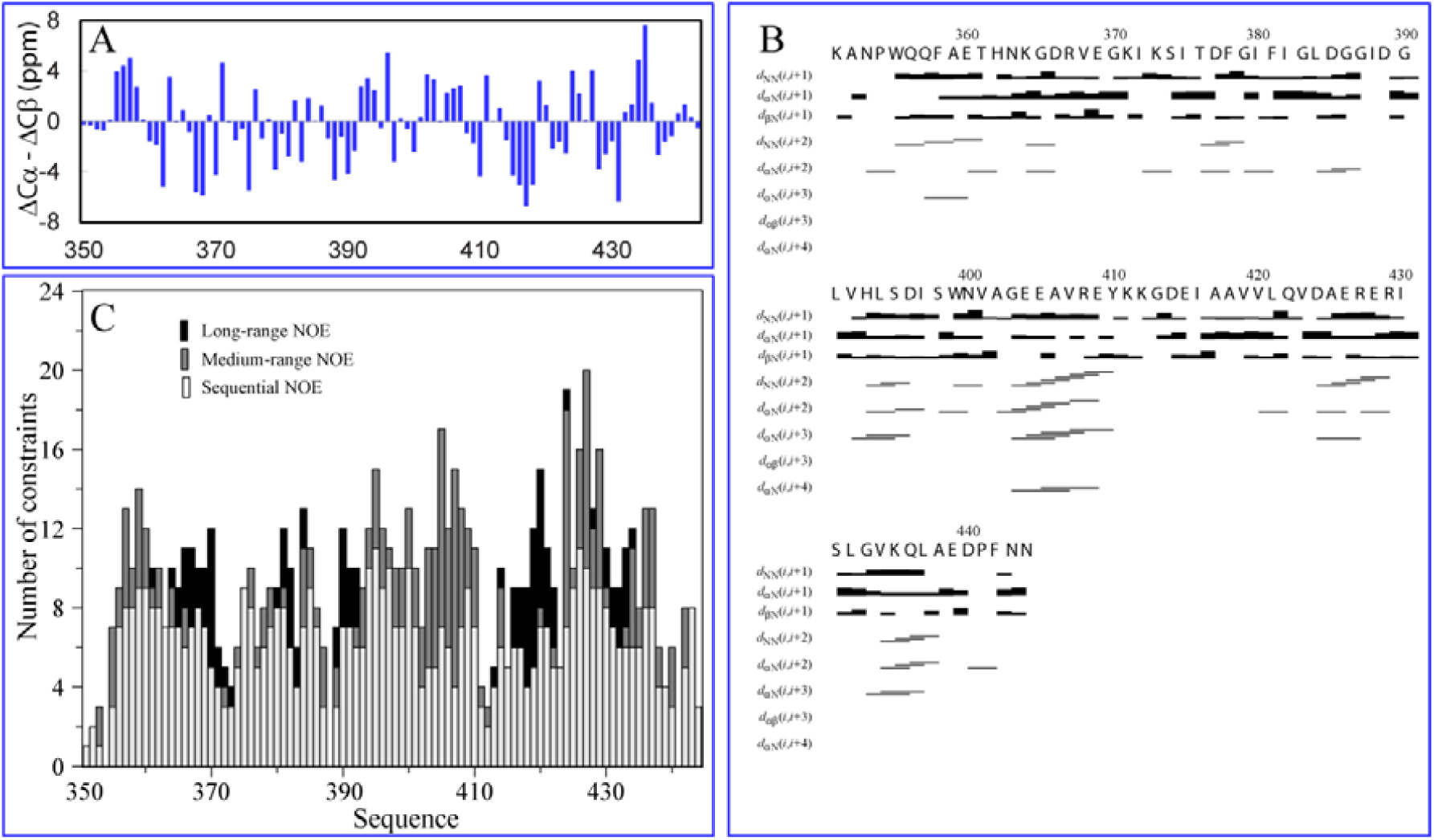
NMR parameters of the D5 domain. (A) Residue-specific (ΔCα-ΔCβ) chemical shifts. (B) NOE patterns defining secondary structures. (C) Residue-specific NOEs categorized into: Sequential, Medium- and Long-range respectively.

**Table S1.** Structural statistics for the 5 lowest energy NMR structures out of 50 calculated for the Domain 5 (residue 350-443) Constraints used for structure calculation

|  |  |
| --- | --- |
| **Constraints used for structure calculation** |  |
| NOE restraints |  |
| Total | 440 |
| Sequential | 288 |
| Medium-range | 97 |
| Long-range $\vert j - i\vert > 5$ | 55 |
| Dihedral angle constraints |  |
| Total | 138 |
| Phi | 69 |
| Psi | 69 |
| Distance violation $> 0.5 \text{ \AA}$ | 0 |
| Angle violation $> 5^\circ$ | 0 |
| Lowest CYANA target function | 3.39 |
| **Deviations from ideal geometry** |  |
| Bond | $0.020 \pm 0.006 \text{ \AA}$ |
| Angles | $2.100 \pm 0.906^\circ$ |
| Improper | $19.550 \pm 7.303^\circ$ |
| **Ramachandran statistics (%)** |  |
| Most favored | 77.5 |
| Additionally allowed | 18.8 |
| Generously allowed | 3.8 |
| Disallowed | 1.3 |
| **Root mean square deviation (RMSD) for all secondary structure fragments ($\text{\AA}$)** |  |
| **(5-12;15-20;30-33;39-42;64-74;78-82;84-87)** |  |
| Backbone atoms | $0.97 \pm 0.18$ |
| Heavy atoms | $1.45 \pm 0.24$ |
| All atoms | $1.92 \pm 0.31$ |
| **Root mean square deviation (RMSD) for the β-barrel regions ($\text{\AA}$)** |  |
| **(15-20;30-33;39-42;64-74;78-82)** |  |
| Backbone atoms | $0.54 \pm 0.11$ |
| Heavy atoms | $1.03 \pm 0.18$ |
| All atoms | $1.41 \pm 0.21$ |

